# Cytotoxicity evaluation of chloroquine and hydroxychloroquine in multiple cell lines and tissues by dynamic imaging system and PBPK model

**DOI:** 10.1101/2020.04.22.056762

**Authors:** Jianling Yang, Meng Wu, Xu Liu, Qi Liu, Zhengyang Guo, Xueting Yao, Yang Liu, Cheng Cui, Haiyan Li, Chunli Song, Dongyang Liu, Lixiang Xue

## Abstract

Chloroquine (CQ) and hydroxychloroquine (HCQ) have been used in treating COVID-19 patients recently. However, both drugs have some contradictions and rare but severe side effects, such as hypoglycemia, retina and cardiac toxicity. To further uncover the toxicity profile of CQ and HCQ in different tissues, we evaluated the cytotoxicity of them in 8 cell lines, and further adopted the physiologically-based pharmacokinetic models (PBPK) to predict the tissue risk respectively. Retina, myocardium, lung, liver, kidney, vascular endothelium and intestinal epithelium originated cells were included in the toxicity evaluation of CQ and HCQ respectively. The proliferation pattern was monitored in 0-72 hours by IncuCyte S3, which could perform long-term continuous image and video of cells upon CQ or HCQ treatment. CC50 and the ratio of tissue trough concentrations to CC50 (R_TTCC_) were brought into predicted toxicity profiles. The CC50 at 24 h, 48 h, 72 h of CQ and HCQ decreased in the time-dependent manner, which indicates the accumulative cytotoxic effect. HCQ was found to be less toxic in 7 cell types except cardiomyocytes H9C2 cells (CC50-48 h=29.55 μM; CC50-72 h=15.26 μM). In addition, R_TTCC_ is significant higher in CQ treatment group compared to HCQ group, which indicates that relative safety of HCQ. Both CQ and HCQ have certain cytotoxicity in time dependent manner which indicates the necessity of short period administration clinically. HCQ has the less impact in 7 cell lines proliferation and less toxicity compared to CQ in heart, liver, kidney and lung.

## Introduction

The Severe Acute Respiratory Syndrome Coronavirus 2 (SARS-CoV-2), was first emerged in China and has spread globally due to its high transmissibility and infectivity, resulting in an unprecedented global public health challenge (1, 2). As of April 20, 2020, more than 2,400,000 cases have been confirmed around the world, according to data supplied by Johns Hopkins University, and at least 58,000 people have died from the disease (2). Judging from current status, most patients have a good prognosis, nevertheless approximately 20% of the patients with COVID-19 experienced critical complications, including arrhythmia, acute kidney injury, pulmonary edema, septic shock, and acute respiratory distress syndrome (ARDS) (3–6). Apart from primarily inflammation in the lungs, it is also suggested that other vital organs like kidneys, heart, gut, as well as liver, were also suffered severe damage according to the autopsies, suggesting that individuals or older with chronic underlying diseases appear to have a higher risk for developing severe outcomes.

Such huge numbers of infected people call for an urgent demand of effective and available drugs to manage the pandemic. Unfortunately, at present, there are still no specific antiviral drugs for prevention or treatment of COVID-19 patients. Recent publications have demonstrated that chloroquine (CQ) and hydroxychloroquine (HCQ) efficiently inhibited SARS-CoV-2 infection *in vitro* assay (7–9). CQ, together with its derivate HCQ, has been commercialized as antimalarial drugs in the clinic for several decades. HCQ has also been broadly used in autoimmune diseases treatment, such as systemic lupus erythematosus (SLE) and rheumatoid arthtitis (10–13). Several clinical trials have confirmed that both CQ and HCQ were superior to the control group in inhibiting the exacerbation of pneumonia, improving lung imaging findings, as well as promoting the virus negative conversion and shorten the disease course.^11^ Moreover, the U.S. Food and Drug Administration (FDA) also approved CQ and HCQ for emergency use to treat hospitalized patients for COVID-19. Although exhibiting apparent efficacy and acceptable safety profile for COVID-19 treatment, CQ and HCQ still have some potential concerns with prolonged usage, including heart rhythm disturbances, gastrointestinal upset, retinal toxicity, in particular for retinopathy(11, 14–17). Additionally, Risambaf et al. found that CQ/HCQ may increase the risk of liver and renal impairment when it used to treat COVID-19(18).

Toxicity tolerability in key tissues about drug effectiveness and side effect were critical to understand their mechanism and to optimize dosing regimen by integrating predicted tissue concentrations (TCs) of both drugs in patients. Therefore, comparison of tissue tolerable concentration and predicted concentration in each given tissue and cell line scan be utilized to suggest dosing optimizing strategy for patients infected by COVID-19, especially in high risk populations. In current study, 8 different types of cell lines including retina, myocardium, lung, liver, kidney, vascular endothelium and intestinal epithelium originated cells were included in the cytotoxic evaluation with the presence of CQ or HCQ in 0-72 h on Incucyte S3 device, which could perform long-term continuous imaging and provide the cellular proliferation pattern upon drug treatment. Consequently, the selectivity index (SI= CC50/TCs) of CQ and HCQ combined with the predicted tissue concentration based on PBPK model was calculated at the given target organ, respectively. The data suggest that HCQ was demonstrated to be much less toxic than CQ, at least at certain key tissues (heart, liver, kidney, and lung). Taken together, this study provides the information regarding cytotoxicity in a wide spectrum and will be beneficial for both pharmacologists and physicians.

## Results

### The effect of CQ on cell proliferation

To gain the more comprehensive cytotoxic information upon CQ and HCQ treatment, we chose 8 different types of cell lines, which included IMR-90, A549, ARPE-19, Hep3B, Vero, HEK-293, H9C2and IEC-6. This panel includes the normal diploid cells, transformed and tumor cell lines which can represent different originated tissue to some extent. To evaluate the cytotoxicity of CQ in the given cell lines list above, we treated them with different dosing regimens of CQ range from 0.017 to 1000 μM. In order to better monitor the effect of CQ on cell viability and proliferation within 0-72 hours, we used the long-term dynamic cell image acquisition device Incucyte S3, which can take photos of cells in each group every three hours. Then the confluence of each group was measured and analyzed by these photos compared with control group. Results from *in vitro* cytotoxicity study showed that CQ exhibited significant cytotoxic at 48 h when the dosing regimens was more than 30 μM. CQ was found to decrease the cell proliferation of in a dose-dependent manner. When the concentration of CQ was more than 300 μM, most of the 8 cell lines showed immediate toxicity within three hours (Figure 1). Among these 8 cell lines, Hep3B, HEK-293, IMR-90, and IEC-6 are more sensitive to CQ.

**Figure 1.**
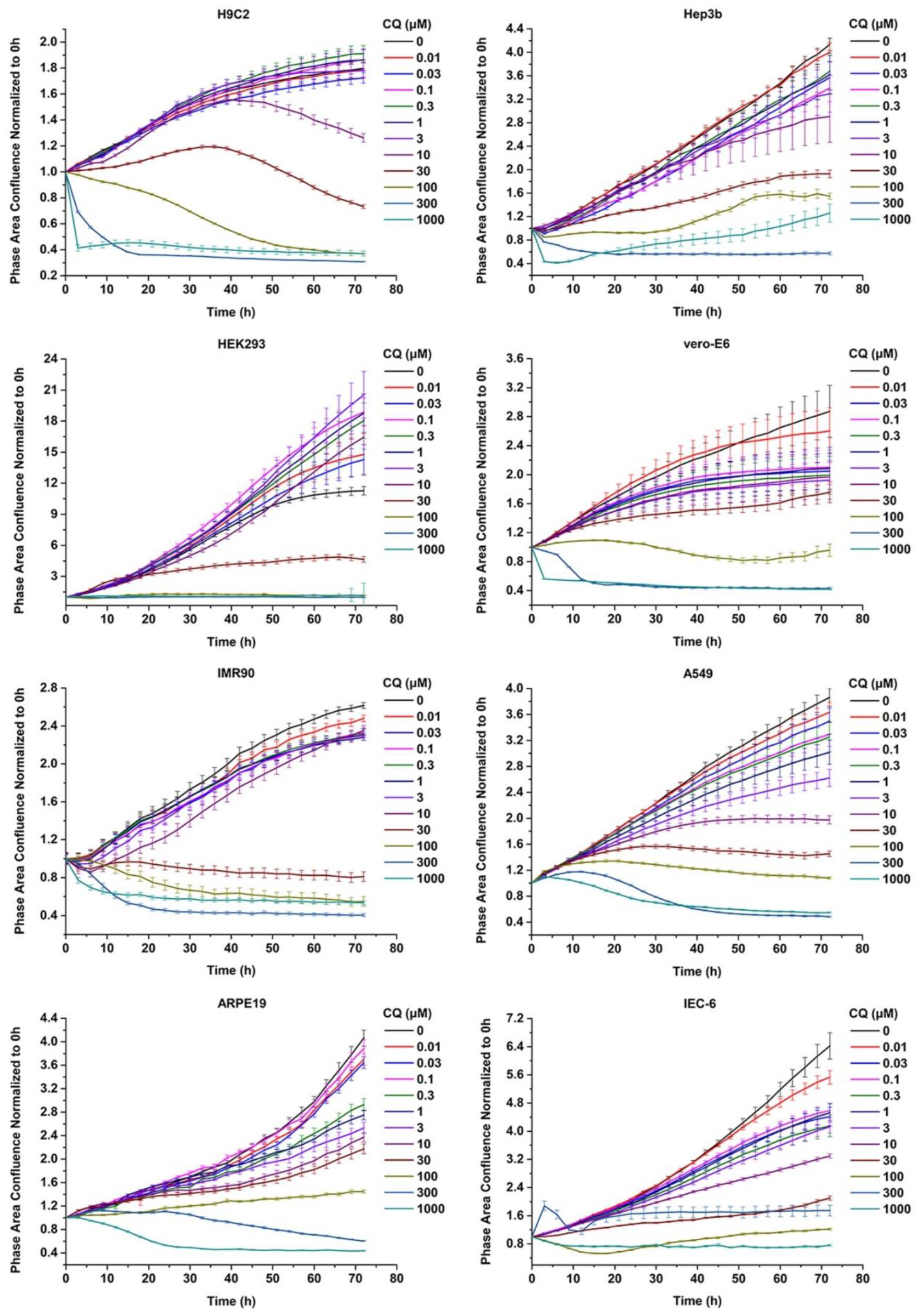
Chloroquine inhibited the viability of the 8 cells in a dose-and timedependent manner. CQ inhibited the viability of Vero E6 cells, IMR90, A549, H9C2, HEK293, Hep3b, ARPE19 cells in a dose-and time-dependent manner. These cells were seeded at a density of 3000-5000 cells per well in a 96-well plate and maintained in regular medium for 72 hours, with different concentration including including 0.01 μM, 0.03 μM, 0.1 μM, 0.3 μM, 1 μM, 3 μM, 10 μM, 30 μM, 100 μM, 300 μM, 1000 μM, respectively. The cell proliferation was assessed by confluence measurements normorlized to 0 hour calculated using IncuCyte (Essen BioScience).The experiments were performed in triplicate.

### The effect of HCQ on cell proliferation

Data from previously reported showed that HCQ also have good antiviral activity for both treatment and pretreatment choice against SARS-CoV-2 (9). So, in the same way as *in vitro* assessment of CQ toxicity, we also test the effect of HCQ on the viability and proliferation of 8 cell lines. Results from the *in vitro* cytotoxicity study showed that HCQ exhibited significant cytotoxic at 48h when the dosing regimens was more than 100 μM. HCQ inhibited the viability of Vero cells, IMR90, A549, H9C2, HEK293, Hep3b and ARPE19 cells in a dose-and time-dependent manner. Among the 8 cell lines, H9C2 and IEC-6 is the most sensitive cell line to HCQ based on the CC50-48 h (Figure 2).

**Figure 2.**
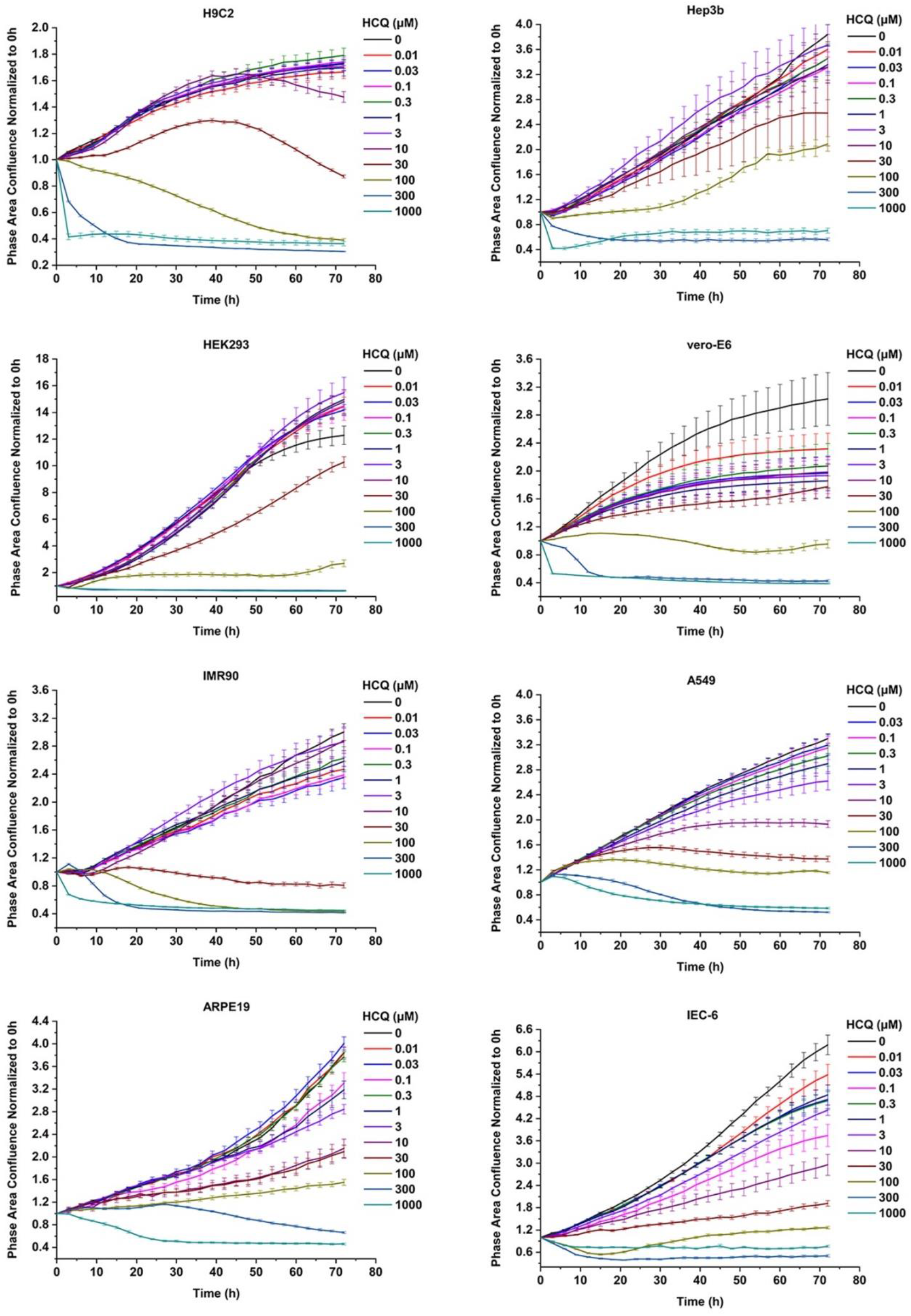
Hydroxychloroquine inhibited the viability of the 8 cells in a dose-and time-dependent manner. HCQ inhibited the viability of Vero, IMR90, A549, H9C2, HEK293, Hep3b, ARPE19 cells in a dose-and time-dependent manner. These cells were seeded at a density of 3000-5000 cells per well in a 96-well plate and maintained in regular medium for 72 hours, with different concentration including 0.01 μM, 0.03 μM, 0.1 μM, 0.3 μM, 1 μM, 3 μM, 10 μM, 30 μM, 100 μM, 300 μM, 1000 μM, respectively. The cell proliferation was assessed by confluence measurements normorlized to 0 Hour calculated using IncuCyte (Essen BioScience).The experiments were performed in triplicate.

### CC_50_ of CQ and HCQ

Cytotoxicity tests were carried out in 8 types of cell lines respectively, which is IMR-90, A549, ARPE-19, Hep3B, Vero, HEK-293, H9C2, and IEC-6 cells and the results are summarized in Table 1 and Figure 3. In this study, CC_50_ values (half cytotoxicity concentration) for CQ and HCQ were measured at 48 h, 72 h respectively. Both CQ and HCQ show strong and immediate toxicity on all 8 cell lines upon treatment more than 300 μM of CQ or HCQ. As shown in Figure 1 and 2, when the concentration of CQ or HCQ is higher than 300 μM, the proliferation shows a sudden decline or brake compared with lower dosing regimens. H9C2 (heart) 、 HEK293(kidney), and IEC-6 (intestine), are the more sensitive cells to CQ compared with 5 other cell lines, as their CC_50_ value at 72 h are less than 20 μM (17.1μM, 16.76 μM, and 17.38 μM respectively). Additionally, the CQ exhibits mild cytotoxic activity on Vero and ARPE-19 cell lines with CC_50_ values of 92.35 μM, and 147.0 μM at 72 h, respectively. Similar with CQ, HCQ exhibits strong cytotoxicity on H9C2 and HEK-293 with CC_50_ values at 72 h lower than 20 μM (15.26 μM and 15.26 μMat 72 h, respectively). HCQ exhibits weak cytotoxic activity on Vero and ARPE-19 cell lines with CC_50_ values of 56.19 μM, and 72.87 μM at 72h, respectively.

**Table 1.** The CC_50_ of CQ and HCQ of 8 cell lines.

| Cell lines | Tissue Type | Drugs | CC50-48h (μM) | CC50-72h (μM) |
| --- | --- | --- | --- | --- |
| H9C2 | Heart | CQ | 41.62 | 17.1 |
|  |  | HCQ | 29.55 | 15.26 |
| Hep3B | Liver | CQ | 36.97 | 24.81 |
|  |  | HCQ | 126 | 108.8 |
| HEK-293 | Kidney | CQ | 16.07 | 16.76 |
|  |  | HCQ | 55.95 | 15.26 |
| Vero | Kidney | CQ | 48.61 | 92.35 |
|  |  | HCQ | 58.22 | 56.19 |
| IMR-90 | Lung | CQ | 29.37 | 25.48 |
|  |  | HCQ | 30.62 | 27.51 |
| A549 | Lung | CQ | 46 | 24.63 |
|  |  | HCQ | 59.86 | 33.05 |
| ARPE-19 | Retina | CQ | 195.4 | 49.24 |
|  |  | HCQ | 208.3 | 72.87 |
| IEC-6 | Intestine | CQ | 32.07 | 23.67 |
|  |  | HCQ | 50.48 | 35.45 |

**Figure 3.**
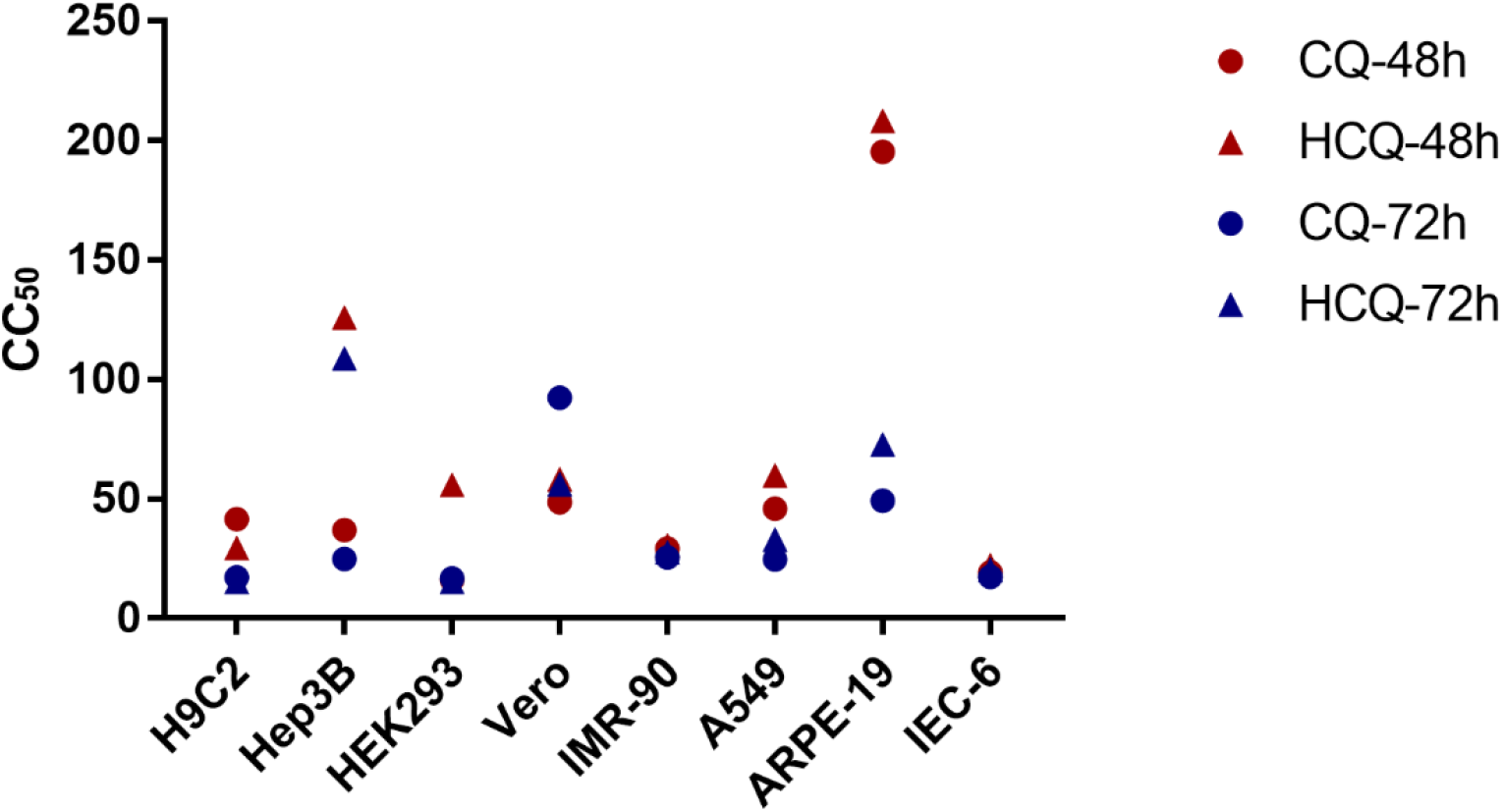
The CC_50_ of CQ and HCQ of 8 different cells in vitro. The CC_50_ (half cytotoxic concentration) of CQ and HCQ were measured by in vitro dynamic imaging system (IncuCyte S3) through monitoring the cell convergence analysis at 0 to 72 h. CC_50_ of CQ and HCQ at 24 h, 48 h, 72 h were analyzed as indicated.

The CC_50_ on 24 h, 48 h, 72 h of CQ and HCQ decreased in a time-dependent manner, which suggests the cumulative toxic effect in most of the 8 cell lines except Vero. As shown in Table 1, the CC_50_ value of 72 h increased instead of decrease compared with that of 48h in Vero, which may be due to special drug metabolism or stability in it. As the selection index (SI) is the safe range to evaluation the drug effect. Considering that the anti-SARS-CoV activity EC50 of HCQ (EC50 = 0.72 μM) is lower than that of CQ (EC50 = 5.47 μM), and the CC_50_ of HCQ is lower than that of CQ in most kinds of cell lines (such as Hep3B, A549, IMR-90, HEK-293 and IEC-6 shown in Table 1) (9). Therefore, we can preliminarily conclude that the selectivity index (SI) of HCQ is higher than that of CQ in most cell types.

### PBPK Model and Risk of Toxicity

Using our PBPK models, we simulated the tissues concentrations of HCQ (600 mg BID for 1 day, 200 mg BID for day 2 to 5) and CQ (500 mg BID for 7 days) (19, 20). The Cmax of tissue concentrations were summarized in Table 2. Results of simulated tissue concentration showed that tissue trough concentration of CQ in liver and lung reached the highest level of drug accumulation (227.545 μg/ml), which is 3 times more than that in heart (60.598 μg /ml). However, the tissue trough concentration of HCQ in lung is the highest level (25.633 μg/ml) compared with liver, kidney and heart (Table 2 and Figure 4).

**Table 2.** Predicted the risk of cytotoxicity in different tissue by R_TTCC_ based on tissue concentration simulated from PBPK model.

| Drug | Value | H9C2 | Hep3B | HEK293 | IMR-90 | Vero | A549 |
| --- | --- | --- | --- | --- | --- | --- | --- |
|  | Tissue | Heart | Liver | Kidney | Lung | Kidney | Lung |
| CQ | PBPK (ng/mL) | 60598.2618 | 227545.0175 | 162787.6437 | 227545.0175 | 162787.6437 | 227545.0175 |
|  | PBPK (μM) | 117.4681 | 441.0898 | 315.5594 | 441.0898 | 315.5594 | 441.0898 |
| | $R_{TTCC}$ | 6.8695 | 17.7787 | 18.8281 | 17.3112 | 3.4170 | 17.9086 |
| HCQ | PBPK (ng/mL) | 6099.0004 | 9546.2615 | 13435.4791 | 25633.4799 | 13435.4791 | 25633.4799 |
|  | PBPK (μM) | 14.0546 | 21.9985 | 30.9609 | 59.0701 | 30.9609 | 59.0701 |
| | $R_{TTCC}$ | 0.9210 | 0.2022 | 2.0289 | 2.1472 | 0.5510 | 1.7873 |
| CQ vs HCQ | $\frac{R_{TTCC}(CQ)}{R_{TTCC}(HCQ)}$ | 7.4586 | 87.9297 | 9.2800 | 8.0621 | 6.2014 | 10.0200 |

**Figure 4.**
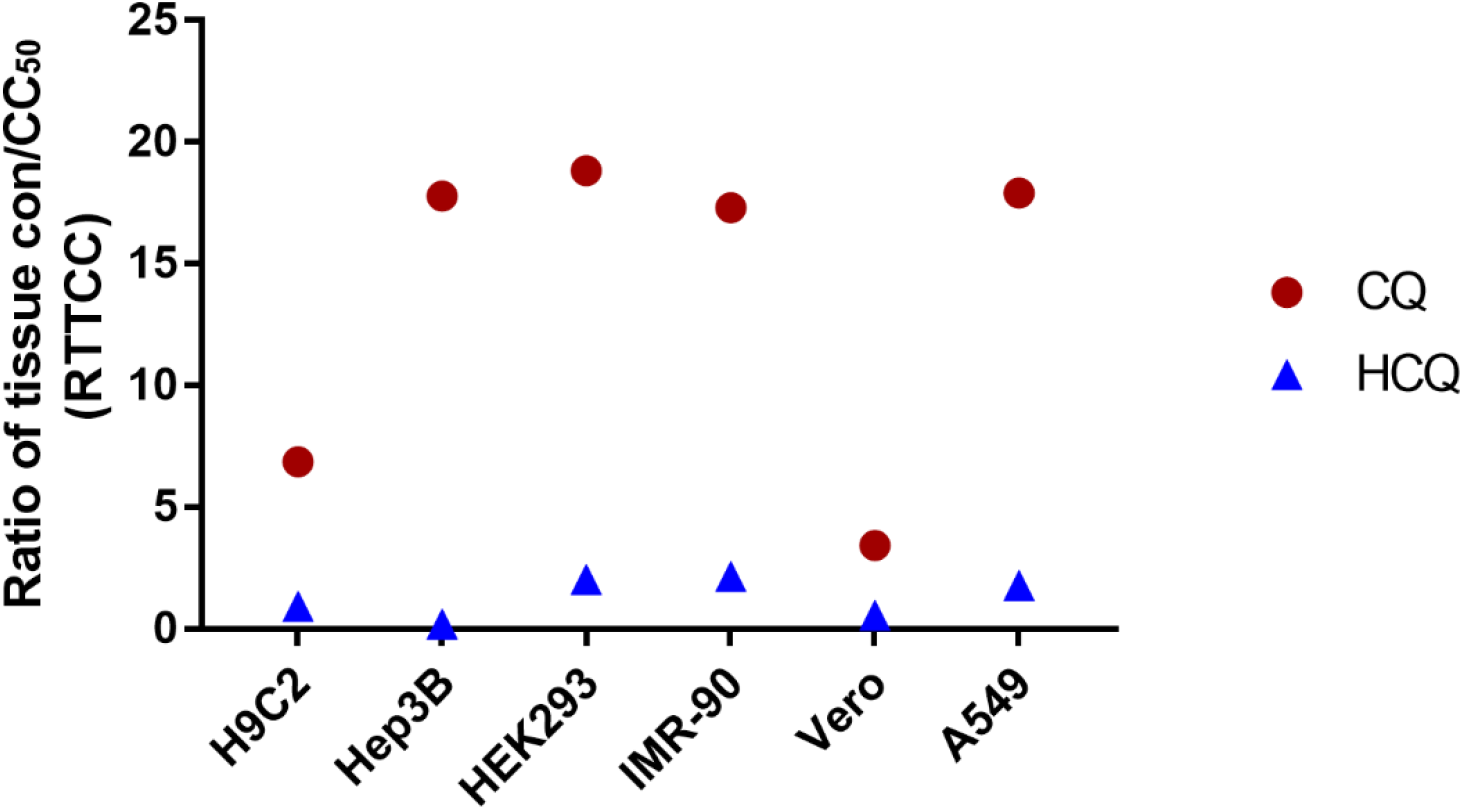
Predicted the risk of cytotoxicity in different tissue by R_TTCC_ based on tissue concentration derived from PBPK model. A. Analysis of ratio of tissue trough concentration vs CC_50_ in 6 cells based on CQ, HCQ tissue concentration simulated by the physiologically-based pharmacokinetic models (PBPK) model by blood data after intravenous administration; B. Compare of R_TTCC_ CQ, HCQ to predict the risk of cytotoxicity in different tissues.

In order to better predict the toxicity risk of CQ and HCQ in different tissues, we used the ratio of simulated tissue trough concentration to CC_50_ (R_TTCC_) to predict the risk of tissue toxicity for the safety profile of these two drugs in the given tissues. As shown in Figure 3, we systematically compared the toxicity between CQ and HCQ, the R_TTCC_ value of CQ is 6-87 times more than that of HCQ in lung, heart, kidney and liver, which suggests that the toxicity risk of HCQ in the above tissues is much lower than that of CQ.

## Discussion

CQ and HCQ, widely-used as antimalarial and autoimmune diseases drugs, recently have been reported that both of them can be used for the treatment of COVID-19 infected patients. As they may block SARS-CoV-2 invasion by inducing the formation of expanded cytoplasmic *in vitro*^(7–9,21, 22)^. In addition, the glycosylation inhibition, together with the pH elevation of endosomes and lysosomes, might be also attributed to their potential antiviral mechanisms (4, 23–25). In addition, the latest findings about HCQ in the application of COVID-19 infected patients suggest that rather than the anti-virus activity, both of them can prevent the cytokine storm by suppressing the immune response (26, 27). Nevertheless, repurposing of CQ or HCQ is an attractive strategy for COVID-19 emergency. Therefore, the potential toxicities of these medications, including gastrointestinal symptoms, cutaneous reactions, cardiotoxicity, hepatotoxicity, in particular retinopathy, are urgent to pay special attention, especially for those elders with underlying diseases.

Our results revealed that both CQ and HCQ have shown certain cytotoxicity in 8 different types of cell lines in time and dose dependent manner *in vitro*, suggesting the necessity of short period administration clinically. Among these types of cell lines, it does show the different tolerant capacity manifested by varied CC_50_ value. For example, the most cytotoxic effect was found in Hep3B (hepatocarcinoma cell line) and IEC-6 (intestinal epithelial cells) treated by CQ, while the A549 (lung cancer), IMR90 (human embryo lung fibroblast cells) and IEC-6 (intestinal epithelial cells) upon HCQ treatment. Although the cytotoxicity was obtained by live cell imaging system *in vitro*, this cellular toxic response of CQ and HCQ may refer to the tissue-toxicity or *vice versa* to some extent. The PBPK models for CQ and HCQ were developed using Simcyp simulator (version 18). Physical and chemical parameters were obtained as previously reported. The lung to blood concentration ratio for CQ and HCQ (obtained from animal studies) was used to predict the drug concentration in the lungs, heart, liver, and kidney. To better investigate the potential toxicity *in vivo* and *in vitro*, we proposed R_TTCC_ (ratio of tissue concentration and CC_50_) derived from PBPK model to predict the risk of toxic profiles in different tissues. We compared the R_TTCC_ data collected from heart, liver, kidney, lung, and revealed HCQ has shown significantly safe profiles than that of upon CQ treatment (9). However, recent publication reported that CQ was safer than HCQ according to SI (7, 9). We speculate that the safety difference might be due to their complex pharmacokinetic characteristics *in vivo*, which possessed specific distribution and long half-life of around days. In short, based on our just published study, we further developed the novel parameters to predict the potential toxicity besides the traditional selectivity index (SI), (the ratio of the CC_50_ to EC_50_), which is a commonly accepted to measure the window between cytotoxicity and antiviral capacity (9). As a result, our data shows that kidney, lung and heart are prone to the toxicity of CQ, otherwise lung and kidney are relative vulnerable upon HCQ treatment (Figure 5). In the meantime, considering the un-negligible effect on cardiocytes and retina cells, of which the most patients with the severe symptoms are more likely suffered the dysfunction in heart and eye sight with aging simultaneously. Therefore, ECG monitoring should be necessary during clinical usage, even for the patients only infected with COVID-19 but without the underlying diseases. In addition, the more attention should be paid to the patients in the changes of their eye sight when using HCQ.

**Figure 5.**
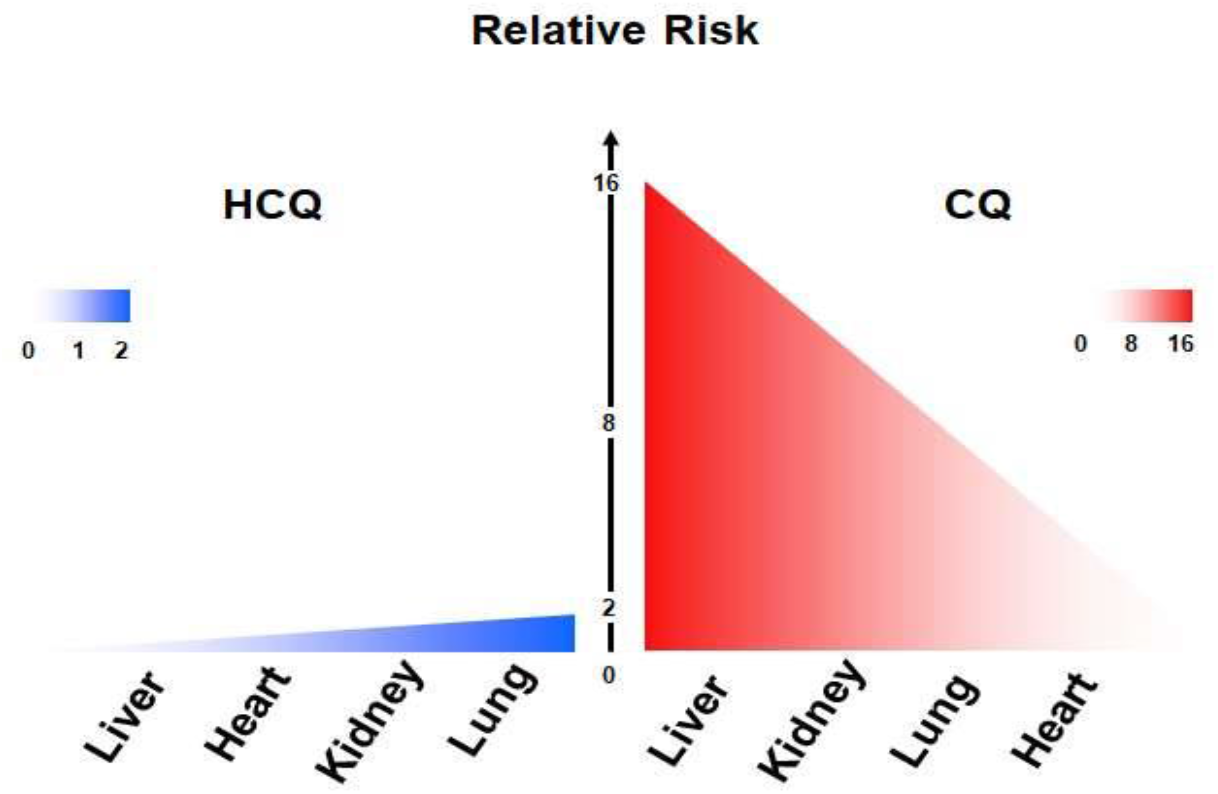
Predicted the risk of toxicity profile for CQ and HCQ. Graphic model for the toxicity of CQ and HCQ in different tissue.

In this study, we perform dynamic imaging system to accurately and precisely monitor the whole proliferation process other than conventional CCK8 assay. Furthermore, R_TTCC_ value suggests that drug distribution should be took in account with the assessment of its potential toxicity within the tissues. Despite of no agreements have been reached on the effectiveness of these candidate drugs in the prevention or treatment of COVID-19, our study could provide more details, new evaluating parameters and deep insight into the safety profile of CQ and HCQ in further preclinical or clinical trials.

## Acknowledgements

We thank Drs Eleanor Howgate and Maurice Dickins for the development of the original chloroquine PBPK base model.

## Abbreviations

SARS-CoV-2: Severe Acute Respiratory Syndrome Coronavirus 2
ARDS: Acute respiratory distress syndrome
CQ: Chloroquine
HCQ: Hydroxychloroquine
COVID-19: Coronavirus disease 2019
SLE: Systemic lupus erythematosus
EC50: Concentration for 50% of maximal effect
CC_50_: The median cytotoxic concentration
PBPK: Physiologically-based pharmacokinetic models
R_TTCC_: Ratio of tissue trough concentration vs CC_50_)

